# A highly efficient method to differentiate CGRP-expressing peptidergic nociceptors from human induced pluripotent stem cells

**DOI:** 10.1101/2025.04.29.651263

**Authors:** Galbha Duggal, Philippa Pettingill, Tatjana Lalic, Shailesh Kumar Gupta, Christine Flodgaard Høgsbro, Viola Volpato, Caleb Webber, Rory Bowden, Despoina Charou, Kanisa Arunasalam, Xinyu Li, Marcello Maresca, Ryan Hicks, Satyan Chintawar, M. Zameel Cader

**Affiliations:** Nuffield Department of Clinical Neurosciences, University of Oxford, Oxford, United Kingdom; Oxford StemTech, The Oxford Science Park, Oxford, United Kingdom; Global Portfolio & Project Management, Oncology R&D, Cambridge UK; Department of Neurology, Faculty of Health and Medical Sciences, Danish Headache Center, Glostrup Hospital, University of Copenhagen, Glostrup, Denmark; UK Dementia Research Institute, Department of Psychological Medicine and Clinical Neuroscience, Cardiff University CF24 4HQ.; Wellcome Trust Centre for Human Genetics, University of Oxford, Oxford, UK; Discovery Biology, Discovery Sciences IMED Biotech Unit, AstraZeneca, Gothenburg, Sweden

## Abstract

Human cellular models of disease developed using induced pluripotent stem cells (iPSC) have enabled wide-ranging research from investigation of human disease mechanisms to phenotypic drug screens. Pain disorders such as neuropathic pain and headache remain areas of considerable unmet need and are considered high risk by pharma. Human iPSC-derived sensory neurons have already been used to accelerate translational research but the current differentiation protocols produce non-peptidergic nociceptors. We demonstrate for the first time the robust differentiation of hiPSC into peptidergic nociceptor lineage with high yield. These nociceptors express CGRP and TRPV1 whilst the non-peptidergic marker RET was completely absent. Functionally, the nociceptors show functional maturity including the expression of TTX-resistant currents and responding to TRPV1 and TRPA1 agonists. Importantly, they were able to release CGRP basally and upon stimulation by an inflammatory soup, which was inhibited upon the application of the 5-HT_1B/1D/1F_ agonist, sumatriptan and topiramate, a migraine prophylactic drug. Overall, we report the successful generation of a novel *in vitro* functional peptidergic nociceptor model which will allow investigation of disease mechanisms in pain and for the development of new effective pain therapies.

---

Sensory neurons can be divided into several categories including nociceptors, mechanoreceptors and proprioceptors based upon the distinct sensory modalities they transduce, such as pain, touch, pressure and body part positions, respectively. Nociceptors are the first-order neurons involved directly in the pathophysiology of pain states including inflammatory pain, neuropathic pain and headache disorders such as migraine. Nociceptors are further divided into myelinated A-δ fibers and unmyelinated/thinly myelinated C-fibers, the latter in turn classified as peptidergic and non-peptidergic nociceptors based upon morphology, molecular markers and functional responses. Specifically, peptidergic nociceptors express neuropeptides such as calcitonin gene-related peptide (CGRP), pituitary adenylate cyclase activating polypeptide-38 (PACAP-38) and Substance P; which are required for thermal nociception and express specific ion channels such as transient receptor potential cation channel subfamily V member 1 (TRPV1, capsaicin receptor) and transient receptor potential cation channel subfamily A member 1 (TRPA1, itch and extreme cold receptor) (Edvinsson et al., 2018a; Liu and Ma, 2011; Wainger et al., 2015). Meanwhile, non-peptidergic nociceptors typically express purinergic receptor P2X3 and transient receptor potential cation channel subfamily M member 8 (TRPM8, cold and menthol receptor) (Liu and Ma, 2011).

Although considerable data has been gathered regarding nociceptor development in rodents and chicks, understanding of human nociceptor development is limited, especially for the peptidergic pathway. Nevertheless, it has been possible using a dual-SMAD inhibition approach to generate functional nociceptors *in vitro* from pluripotent stem cells including human embryonic stem cells (hESCs) and human induced pluripotent stem cells (hiPSCs) (Chambers et al., 2012). These nociceptors expressed RUNX1 and responded to ATP indicating a non-peptidergic DRG lineage (Chambers et al., 2012), with minimal response to capsaicin despite prominent TRPV1 expression (Röderer et al., 2023; Schoepf et al., 2020; Young et al., 2014). A similar dual-SMAD inhibition strategy was adopted to generate trigeminal ganglion (TG)-nociceptors from hESCs(Dincer et al., 2013), however, SMAD1/5/8 inhibition was removed at day 3 in order to induce a cranial placodal fate. More recent attempts on peptidergic cell fates focused on TRPV1 responsiveness, either via chemical induction (Deng et al., 2023; Guimarães et al., 2018; Muller et al., 2018; Wilson et al., 2018), ectopic expression of key transcription factors (Blanchard et al., 2015; Wainger et al., 2015), or a combination of both (Hulme et al., 2020; Lee et al., 2015; Schrenk-Siemens et al., 2022; Vojnits et al., 2019). Starting materials were not limited to pluripotent stem cells, also including fibroblasts, epidermal neural crest stem cells, or blood iNPCs. However, only a few tested for secretive neuropeptides, e.g. Substance P (Deng et al., 2023; Guimarães et al., 2018; Muller et al., 2018), and CGRP (Muller et al., 2018) ELISA in human cells. Although promising CGRP release was shown in murine fibroblast-devired nociceptors, the overall yield of peptidergic neurons was low and nociceptors derived from human fibroblasts were not investigated for CGRP expression and release (Wainger et al., 2015).

Peptidergic nociceptors are of considerable interest in disorders such as migraine where the neuropeptide CGRP has been demonstrated to play a prominent role in its pathophysiology (Edvinsson and Warfvinge, 2019). Upon activation, trigeminal primary afferents release CGRP to cause neurogenic inflammation and vasodilation (Russell et al., 2014). Hence, peptidergic nociceptors regulate nociception, modulate peripheral and cerebral blood flow and initiate or sustain inflammatory processes (Benemei et al., 2017; Lukacs et al., 2017). Furthermore, CGRP seems to be elevated during a migraine attack and CGRP receptor antagonists (Bell, 2014) and more recently CGRP monoclonal antibodies (Goadsby et al., 2017; Oliveira et al., 2024; Silberstein et al., 2017) are showing good efficacy in migraine clinical trials, with the first migraine preventative drug recently approved by the U.S. Food and Drug Administration (Edvinsson et al., 2018b).

As the current established protocols for nociceptor differentiation recapitulate only the neural crest non-peptidergic fate *in vitro*, we aimed to formulate an optimized protocol using hiPSCs to induce the peptidergic nociceptive lineage. Here, we demonstrate a method which can be adapted to produce either non-peptidergic TRPA1-expressing nociceptors (∼45%) which has not been shown previously or TRPV1/CGRP expressing peptidergic nociceptors (∼93%). The peptidergic nociceptors were able to release CGRP and responded to nociceptive stimuli including capsaicin and inflammatory soup or efficacious migraine therapies. Remarkably, we show that by co-culture with astrocytes, *in vitro*-derived TG early nociceptors can switch their fate from a non-peptidergic to peptidergic nociceptor type. The highly efficient production of peptidergic nociceptors *in vitro* from hiPSCs opens new avenues in drug-disease modelling thereby enhancing the possibilities of successful therapies for pain.

## Results

### Optimization and temporal molecular profiling of human iPSC-derived TG nociceptors for peptidergic and non-peptidergic nociceptive markers

To understand the *in vitro* specification of nociceptors towards a peptidergic or a non-peptidergic fate, we adopted the TG-nociceptor specific differentiation protocol as previously described (Dincer et al., 2013) with culture modifications from the placodal isolation stage (Fig. 1a). This involved dual-SMAD inhibition of human iPSCs followed by placode induction and we observed successful switching of developmental fates from neuroectoderm to pre-placodal lineage, as exhibited by downregulation of PAX6 and upregulation of SIX1 (Fig. 1b). We found the nociceptors generated using the established protocol were functionally immature when cultured on PO/LAM coating and were CASPASE3-positive, thereby suggesting compromised cell survival in these cultures (Supplementary Fig. 1a). In order to circumvent this issue, we replaced the substrate from PO/LAM to matrigel and supplemented forskolin (FK) in our maturation growth media. We chose matrigel as it has been shown to enhance neuronal survival and maturation due to its composition of high levels of brain extracellular matrix proteins (Choi et al., 2014; Uemura et al., 2010). Additionally, FK, a protein kinase A agonist, elevates cyclic AMP (cAMP) which leads to phosphorylation and inactivation of GSK3β thereby repressing the apoptotic activity in neurons. FK is also known to induce the expression of voltage-gated ion channels, formation of functional synaptic contacts and facilitate neuronal functional maturation in neural progenitor cells (Lepski et al., 2013; Li et al., 2000). We found that the combination of matrigel and FK upon placodal isolation generated functionally mature neurons displaying a physiological resting membrane potential of -60mV and were CASPASE3-negative (Supplementary Fig. 1). Next, to determine the timing of divergence of nociceptive population towards the different fates, we conducted RUNX1/cMET expression profiling at the key stages of differentiation. We found that as differentiation progresses, the expression of non-peptidergic marker RUNX1 significantly increases whereas the early peptidergic marker cMET exhibits a reverse profile, both at the transcript and protein level (Fig. 2a). *CGRP* expression also appeared to match the *cMET* expression profile, with increased level at the post-mitotic placodal stage (days 13-17) followed by significant downregulation (Fig. 2a). Interestingly, there were rare populations (∼1-2%) of RUNX1^-^/cMET^+^ cells as observed by immunocytochemistry (Fig. 2a), which may indicate a potential of these cells to generate peptidergic nociceptors or perhaps a distinct subclass of nociceptive neurons. Similar to TG differentiation, the expression profiling of RUNX1/cMET during DRG differentiation revealed increase in non-peptidergic marker RUNX1 and downregulation of peptidergic markers cMET and CGRP, matching with *in utero* observation of mouse embryo (Chen et al., 2006; Kobayashi et al., 2012) both at transcript and protein level as differentiation progressed (Supplementary Fig. 2). However, unlike TG differentiation, which displayed pockets of peptidergic-like RUNX1^-^/cMET^+^ cells, DRG differentiation generated regions of non-peptidergic RUNX1^+^/cMET^-^ expressing cells (Supplementary Fig. 2) suggesting their decreased *in vitro* differentiation propensity towards a peptidergic lineage.

**Figure 1:**
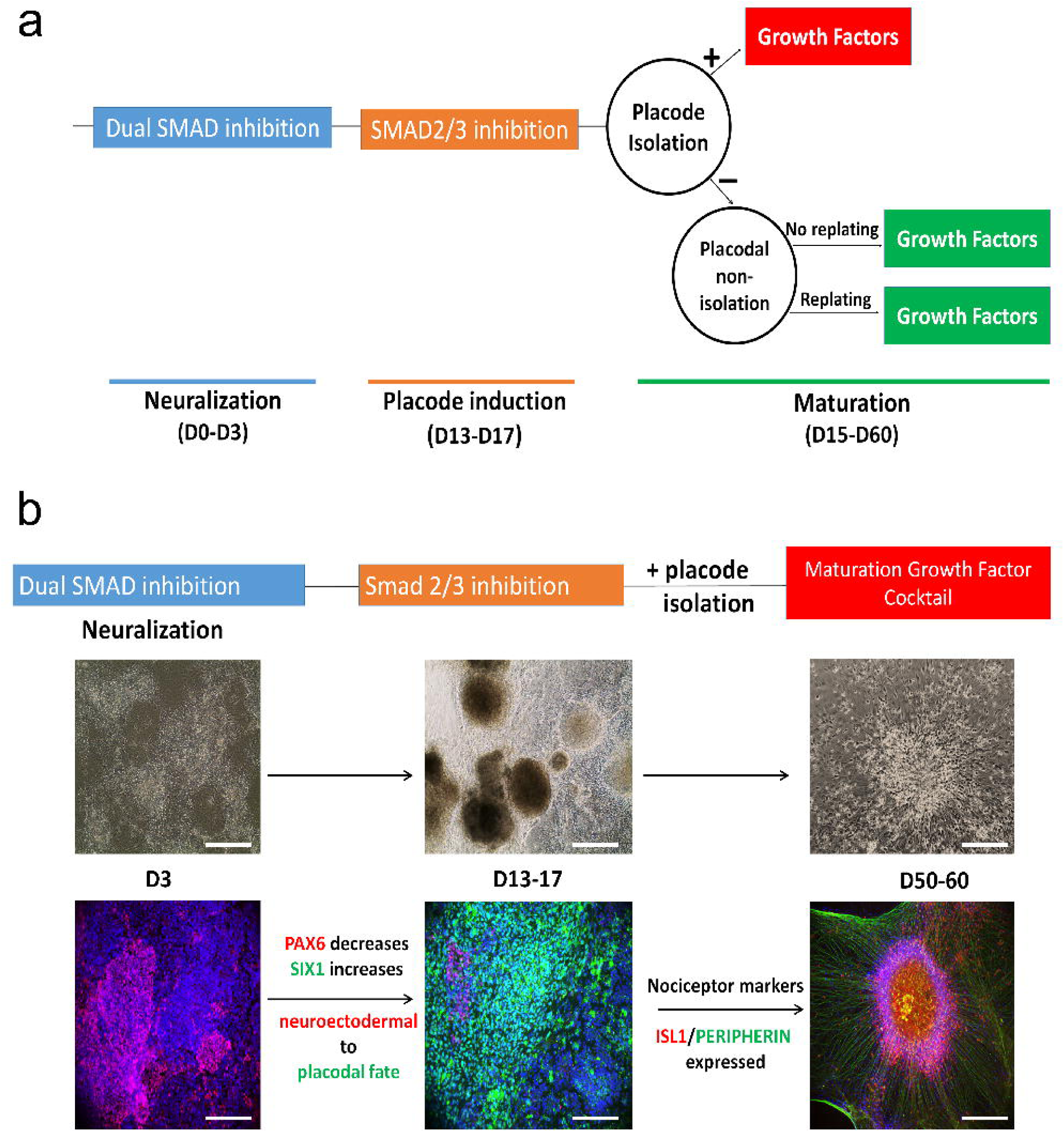
Differentiation of human iPSCs towards TG nociceptors. **a)** Schematic representation of modified TG nociceptor differentiation protocol. Human iPSCs are subjected to dual SMAD inhibition and either subjected to placodal isolation, or no isolation or no isolation and replating strategies, and undergo maturation until day 60. **b)** Validation of directed differentiation of human iPSCs towards TG nociceptive fate, whereby cranial placode-specific SIX1 marker is expressed followed by the expression of sensory neuron markers ISL1 and PERIPHERIN upon maturation, scale bar 10µm.

**Figure 2:**
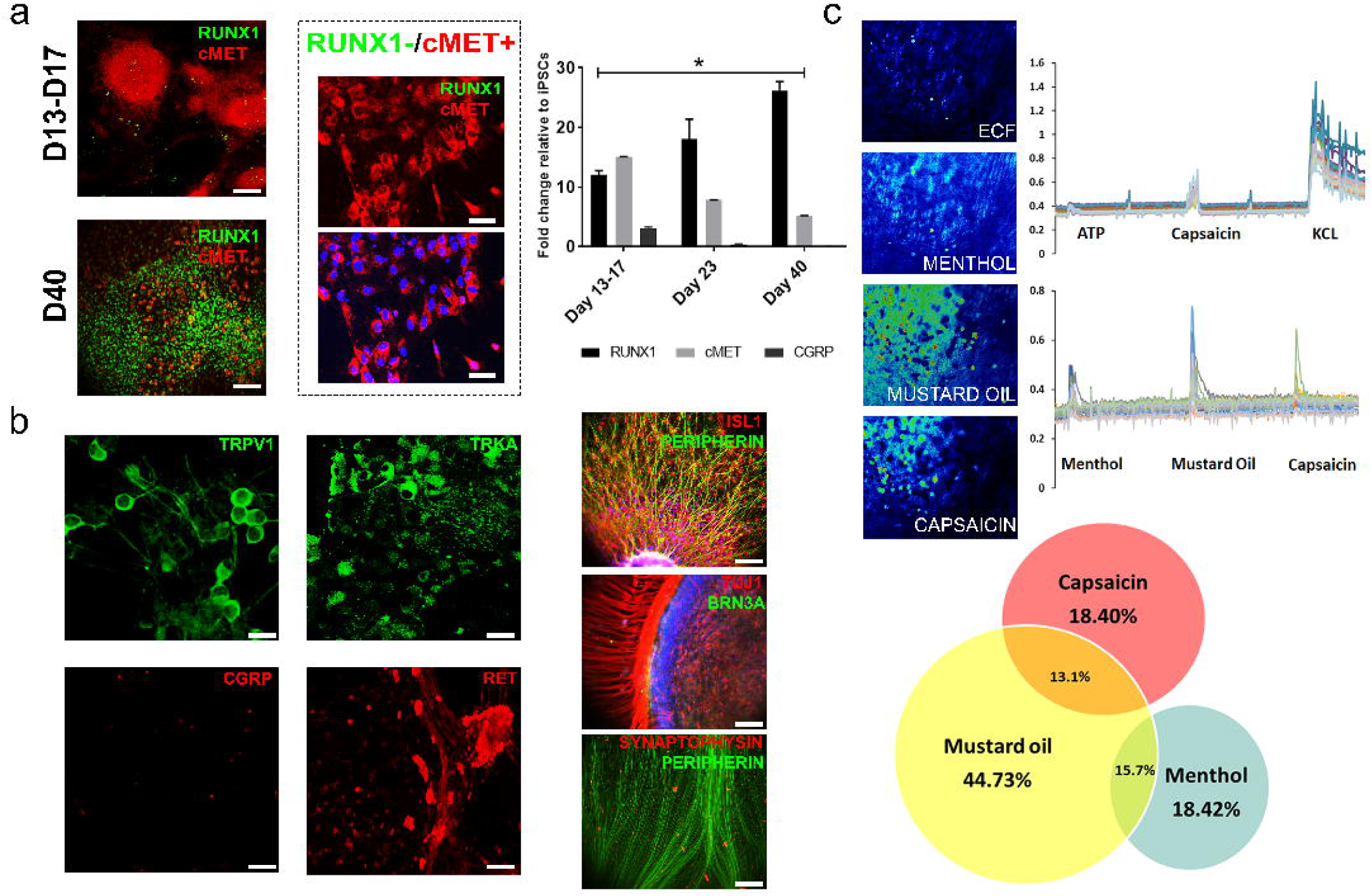
TG placodal isolation results in the generation of RUNX1^+^/RET^+^/TRKA^+^/ TRPV1^+^/TRPA1-responsive non-peptidergic nociceptor subtypes. **a)** (Left panel) RUNX1 expression increases while cMET expression decreases from post-mitotic placodal stage to day 40 of neuronal maturation, scale bar 10µm. Dashed box reveals small population that has RUNX1^-^/cMET^+^ peptidergic identity at day 40, scale bar 25µm. (Right panel) Gene expression profiling by qRT-PCR also reveals significant increase in non-peptidergic *RUNX1* marker while peptidergic-specific *cMET* and *CGRP* marker expression is significantly reduced (p<0.05, n = 3 from three independent iPSC differentiation rounds). **b)** (Left panel) Immunocytochemistry reveals that day 60 mature TG nociceptors express TRPV1, TRKA and RET while CGRP continues to be lowly expressed, scale bar 50µm. (Right panel) Validation by immunocytochemistry for nociceptive markers PERIPHERIN, ISL1, BRN3A, neuronal marker TUJ1 and mature neuronal marker SYNAPTOPHYSIN which further confirms their mature sensory neuronal identity, scale bar 10µm. **c)** Calcium imaging demonstrates that TG-placodal isolated nociceptors respond robustly to TRPA1 agonist mustard oil followed by high dose of TRPV1-agonist capsaicin giving them a non-peptidergic identity (n = 67 cells, pooled from three independent rounds of iPSC differentiation).

### TG placodal isolation promotes the generation of TRPV1/TRPA1 non-peptidergic nociceptors

We next characterized the identity of the matured *in vitro*-derived TG nociceptors generated upon placodal isolation in our optimized culture conditions. Interestingly, these TG-nociceptors expressed TRPV1 and TRKA while CGRP expression was almost absent (Fig. 2b). Immunocytochemistry revealed that the nociceptors continue to express the non-peptidergic marker RET (Fig. 2b). We also confirmed the expression of PERIPHERIN, ISL1, BRN3A, TUJ1 and SYNAPTOPHYSIN (Fig. 2b). Sensory neurons *in vivo* display a varied response to different TRP channel agonists (Levine and Alessandri-Haber, 2007; Usoskin et al., 2015), and we performed an evaluation of calcium flux in these TG-nociceptors to determine their functional competence. For this purpose, a series of agonists were utilized to detect the ion channel and receptors which respond to noxious stimuli including P2X3 (10µM ATP), TRPM8 (100nM menthol), TRPA1 (250µM allyl isothiocyanate or mustard oil) and TRPV1 (25µM capsaicin) and the number of responders were quantified with a stable baseline and as a percentage of KCl-responders (Fig. 2c). Although we observed rapid calcium fluctuations in response to menthol, mustard oil and capsaicin, we found that majority of the neurons (44.73%) belonged to mustard oil-responsive TRPA1 nociceptive lineage, 18.40% exhibited activation of TRPV1 channels upon exposure to capsaicin and of these, 13% responded to both mustard oil and capsaicin. 18.4% of neurons responded to menthol and 15% of these responded to both mustard oil and menthol. We found very few cells responded to ATP, and no cells responded to both capsaicin and menthol (Fig. 3c, n = 67 cells). The lack of ATP response is consistent with rodent TG neurons, where most IB4 non-peptidergic neurons do not express ATP-receptor P2X3, unlike DRG IB4 non-peptidergic neurons which do express P2X3 (Ambalavanar et al., 2005). Therefore, our finding is consistent with the TG origin of our non-peptidergic nociceptors.

**Figure 3:**
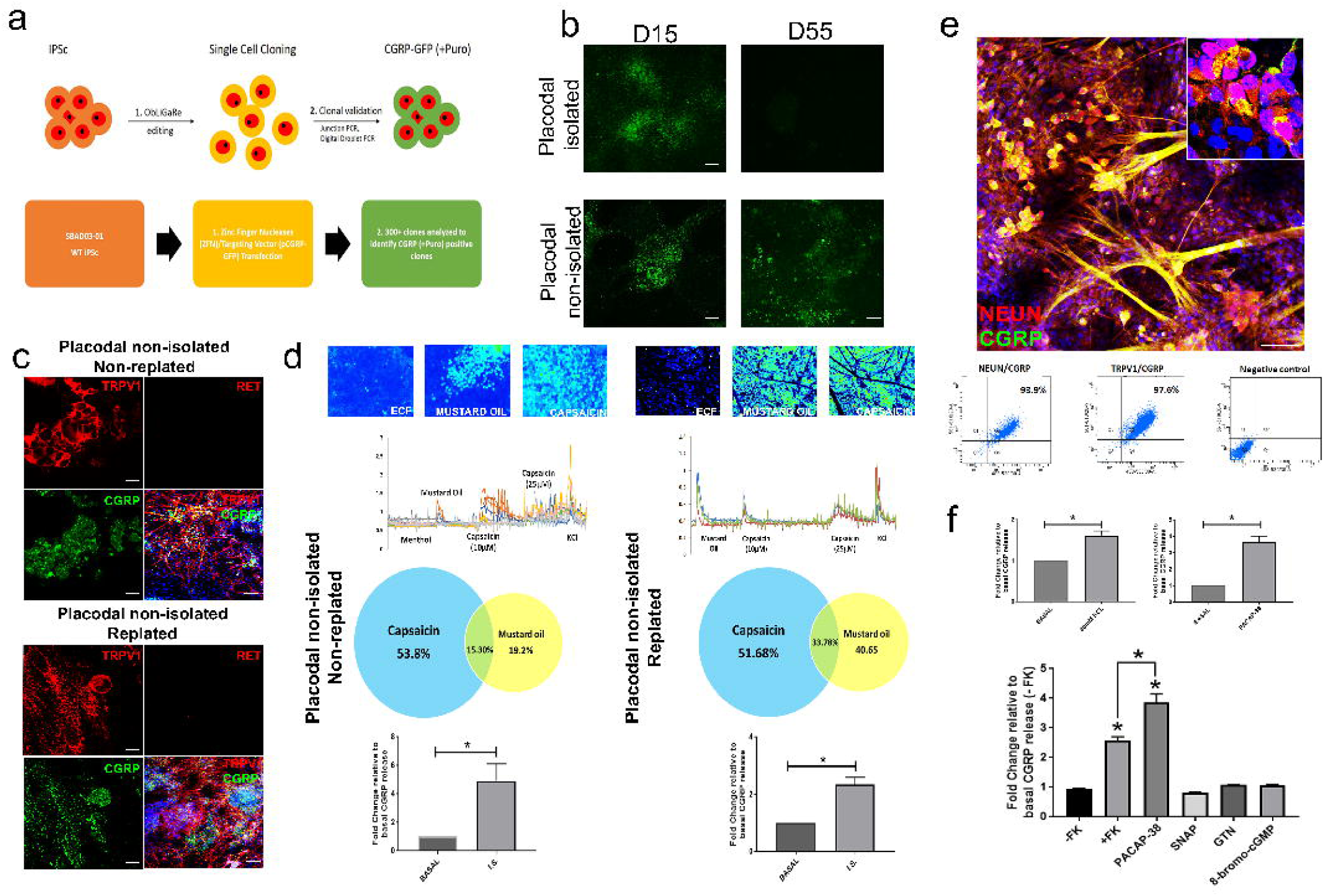
TG nociceptors derived in the absence of placodal isolation (± replating) release CGRP similar to peptidergic nociceptors. **a)** Workflow for generation of CGRP-GFP reporter line. **b)** CGRP-GFP expressed in mature TG nociceptors in the absence of placodal isolation, with or without replating (bottom panel, scale bar 5µm). **c)** Immunocytochemistry reveals co-expression of CGRP and TRPV1 while RET expression is absent suggesting peptidergic subtypes, scale bars 5µm, 50µm. **d)** Functional analysis by calcium imaging and ELISA on non-GFP iPSC lines, for placodal non-isolated non-replated (left panel, n = 54 cells) placodal non-isolated replated (right panel, n = 35 cells). Calcium imaging shows that these nociceptors are TRPA1/TRPV1-responsive to low dose capsaicin, characteristic of peptidergic sensory neurons. Three independent rounds of differentiation were performed. They also undergo significant CGRP release upon stimulation by inflammatory soup (I.S.) (n = 3 dishes per stimulation per differentiation round, fold change quantified relative to basal release per stimulation, p<0.05). **e)** Immunocytochemistry and analysis by flow cytometry for placodal non-isolated replated cultures demonstrates co-localization of mature neuronal marker NEUN with peptidergic marker CGRP, further validating their nociceptive identity, scale bars 5µm, 50µm. **f)** (Top panel) CGRP release quantified by ELISA upon stimulation with 80mM KCl and 1µM PACAP-38. (Bottom panel) CGRP release in response to cAMP (FK, PACAP-38) and cGMP (SNAP, GTN, 8-bromo-cGMP) agonists (n = 3 dishes per stimulation per differentiation round, fold change quantified relative to basal release per stimulation, p<0.05)

### Generation of functionally competent *in vitro*-derived TG capsaicin-responsive peptidergic nociceptors

In order to better understand the timing of CGRP expression during nociceptor-differentiation, we generated CGRP (CALCA gene promoter)-GFP knock in reporter lines by targeting the *AAVS1* locus into human chromosome 19 (in the intron one of *PPP1R12C* gene) using Zinc Finger Nucleases (ZFNs) mediated gene editing method known as ObLiGaRe (Maresca et al., 2013) (Obligate Ligation-Gated Recombination) (Fig. 3a, Supplementary Fig. 3, 4). Using the CGRP-GFP reporter iPSCs we found that consistent with our *CGRP* expression analysis, CGRP-GFP reporter was robustly present up to day 15 in the placodal stage but was extinguished shortly after placodal isolation and remained silent in the mature cultures (Fig. 3b). Whereas, in the absence of placodal isolation, CGRP-GFP expression was maintained in the mature day 55 cultures, both in non-replated and replated cultures (Fig. 3b). We therefore compared the two culture conditions (placodal non-isolated ± replating) to determine whether leaving placodes with the surrounding cells might aid in the generation of peptidergic neurons upon maturation.

Using immunofluorescence, we confirmed the placodal non-isolated TG cultures (± replating) co-expressed TRPV1/CGRP, whilst RET expression was absent (Fig. 3c). Functionally, the placodal non-isolated non-replated TG cultures were responsive to capsaicin and they appeared to group into cells with increased receptor sensitivity to 10µM capsaicin (53.8%) and cells only responding to a higher dose, 25uM capsaicin (23%) (Fig. 3d). Some neurons also responded to mustard oil (19.2%) and approximately 15.3% nociceptors responded to both capsaicin and mustard oil (Fig. 3d, n = 54 cells). Functional analysis of the placodal non-isolated replated nociceptors by calcium imaging again confirmed responsiveness to 10uM capsaicin (51.68%). Interestingly we found a greater proportion of cells responding to mustard oil (40.65%) and 33.78% nociceptors were responsive to both capsaicin and mustard oil (Fig. 3d, n = 35 cells). There were no cells that responded to menthol in either of the culture strategies.

Although we were able to generate functional peptidergic nociceptive population, the placodal non-isolated non-replated cultures were extremely confluent impeding their use in downstream assays including electrophysiology. Therefore, adopting the replating strategy enabled peptidergic neuronal culture expansion for several downstream analyses in a cost-effective manner. Importantly, using our replating method, we were able to isolate neurons as single cells at day 55 for quantification by flow cytometry, which revealed that 97.6% cells were TRPV1/CGRP-positive and 93.9% cells co-expressed NEUN and CGRP (Fig. 3e). This therefore confirms that our protocol generates a highly pure iPSC-derived TRPV1/CGRP TG-peptidergic population. The mature peptidergic nociceptive identity of these replated cultures was further confirmed by the expression of ISL1, PERIPHERIN, TUJ1 and SYNAPTOPHYSIN (Supplementary Fig. 5)

To confirm the functional maturation of the iPSC-derived peptidergic nociceptors using the replating method, we performed whole-cell patch clamp recordings to determine their electrophysiological properties. Trains of action potentials were elicited in response to depolarizing current injection in current-clamp recordings. On some occasions spontaneous synaptic activity could be observed (Supplementary Fig. 6). Voltage clamp recordings were made to further demonstrate that iPSC-derived neurons express the repertoire of voltage-gated ion channels characteristic of functional neurons. All recorded cells (n = 15) showed fast voltage activated inward currents followed by slow outward currents, consistent with sodium and potassium currents, respectively (Supplementary Fig. 6). Inward currents were blocked by 1µM tetrodotoxin (TTX), a sodium voltage-gated channel blocker and TTX-resistant currents were also observed (Supplementary Fig. 6).

### Modulation of CGRP release by migraine provocants and migraine treatments

To examine CGRP release, we chose stimulation by inflammatory soup, composed of ATP, bradykinin, PGE_2_, histamine and noradrenaline, as it is known to potently facilitate CGRP release in TG rodent primary cultures (Basbaum et al., 2009; Durham and Russo, 1999). We observed that exposure to inflammatory soup resulted in 5-fold and 3-fold increase in CGRP release compared to basal levels both in non-replated and replated cultures, respectively (Maingret et al., 2008) (Fig. 3d, n = 3 separate dishes per iPSC line).

We next assessed CGRP release in response to depolarization signal by KCl as well as to the migraine provocants: neuropeptide PACAP-38 (Vollesen and Ashina, 2017) (Fig. 3f), nitroglycerin (GTN) and sodium nitroprusside (SNAP). CGRP release was significantly elevated by 1.5-fold in response 80mM KCl stimulation (p<0.05), while PACAP-38 induced 4-fold increase in CGRP levels (p<0.05) compared to the basal release (n = 3 separate dishes, Fig. 3f). CGRP release was also provoked by FK but no release was observed with SNAP, GTN or 8-boromo-cGMP.

Many of the current migraine drugs may mediate their effects through altering CGRP release and hence, we examined the acute migraine drug, sumatriptan and the migraine preventative, topiramate. We found that inflammatory soup evoked CGRP release was significantly inhibited in the presence of sumatriptan and topiramate (n = 3 separate dishes per iPSC line, Fig. 4) and we observed the expression of 5-HT_1D_ receptor (site of action of Sumatriptan) in these replated TG peptidergic cultures (Supplementary Fig. 7).

**Figure 4:**
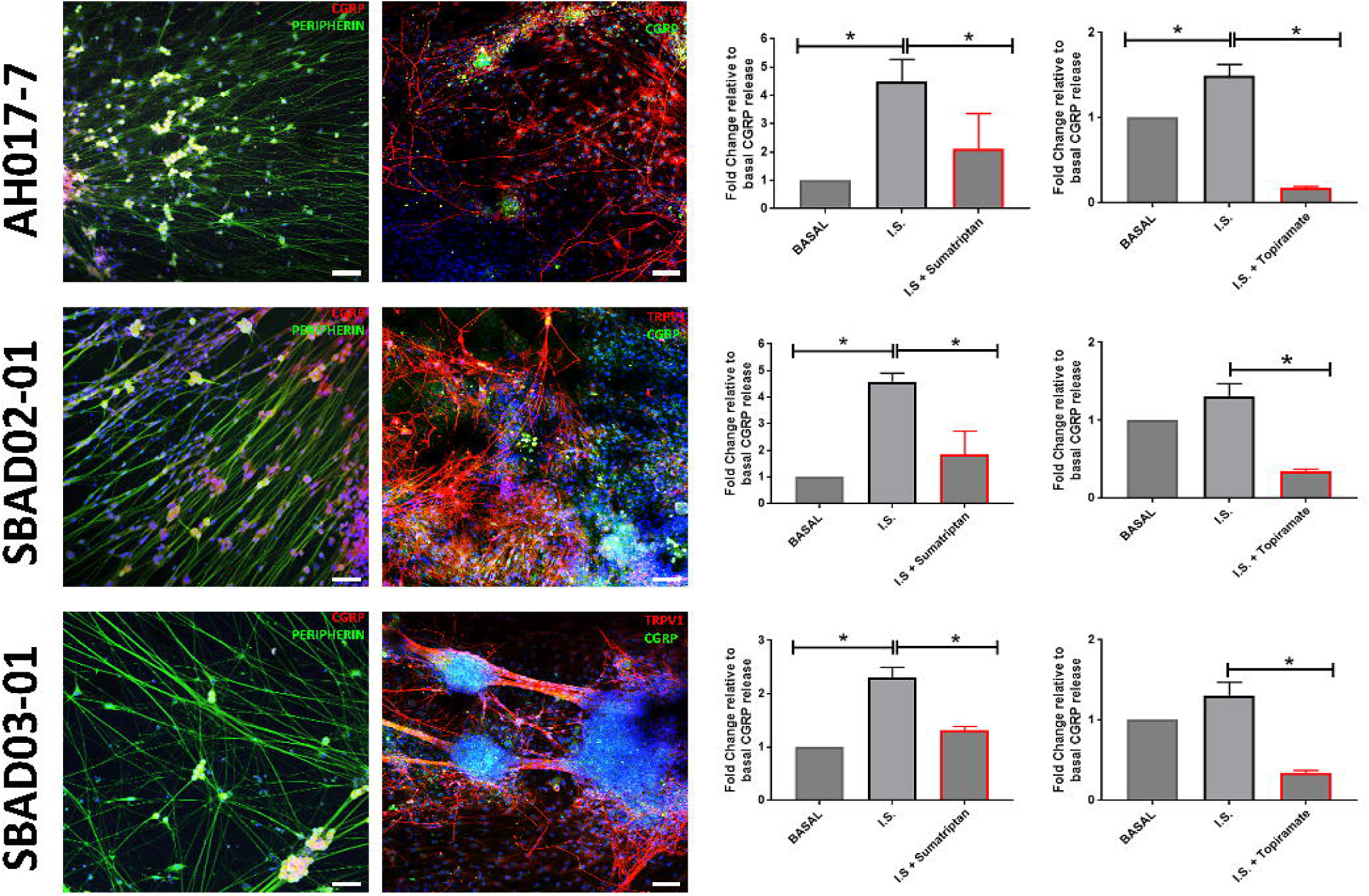
Efficient reproducibility of peptidergic differentiation protocol in AH017-7, SBAD03-01 and SBAD02-01 human iPSCs lines. (Left panel) Immunocytochemistry reveals peptidergic nociceptor identity of the TG neurons as confirmed by the expression of CGRP, PERIPHERIN and TRPV1. (Right panel) Drug-inhibition assay on I.S.-stimulated CGRP release quantified by ELISA (p<0.05, n = 3 separate dishes per iPSC line per stimulation, per differentiation for each iPSC line, fold change quantified relative to basal release per stimulation, p<0.05).

### Single cell transcriptomic analysis reveals human TG-like characteristics of the *in vitro*-derived TG peptidergic model

To confirm the identity of iPSC-derived replated TG nociceptor cultures, we undertook single cell transcriptomics using the 10X genomics platform. We constructed a cell type-specific gene expression space through principal component analysis of different human brain cell types and then projected iPSC-derived nociceptor single cells from the control samples onto this space. We observed that all iPSC-derived nociceptor cells clustered much closer to the external TG/DRG samples than to the other cell types (Flegel et al., 2015; LaPaglia et al., 2018) (Fig. 5). Moreover, while these iPSC-derived nociceptors could be formed into distinct clusters based upon their transcriptomic variation, once projected onto the cell type-specific gene expression space, all iPSC-derived nociceptor cells did not separate by the identified clusters suggesting cell clusters were not defined by varying cell identity (Fig. 5). For this reason and because of the very low and noisy expression of marker genes at the single cell level, we merged the reads from all cells to create a pseudo-bulk sample that we further compared with external TG/DRG samples based on selected marker genes (Fig. 5).

**Figure 5:**
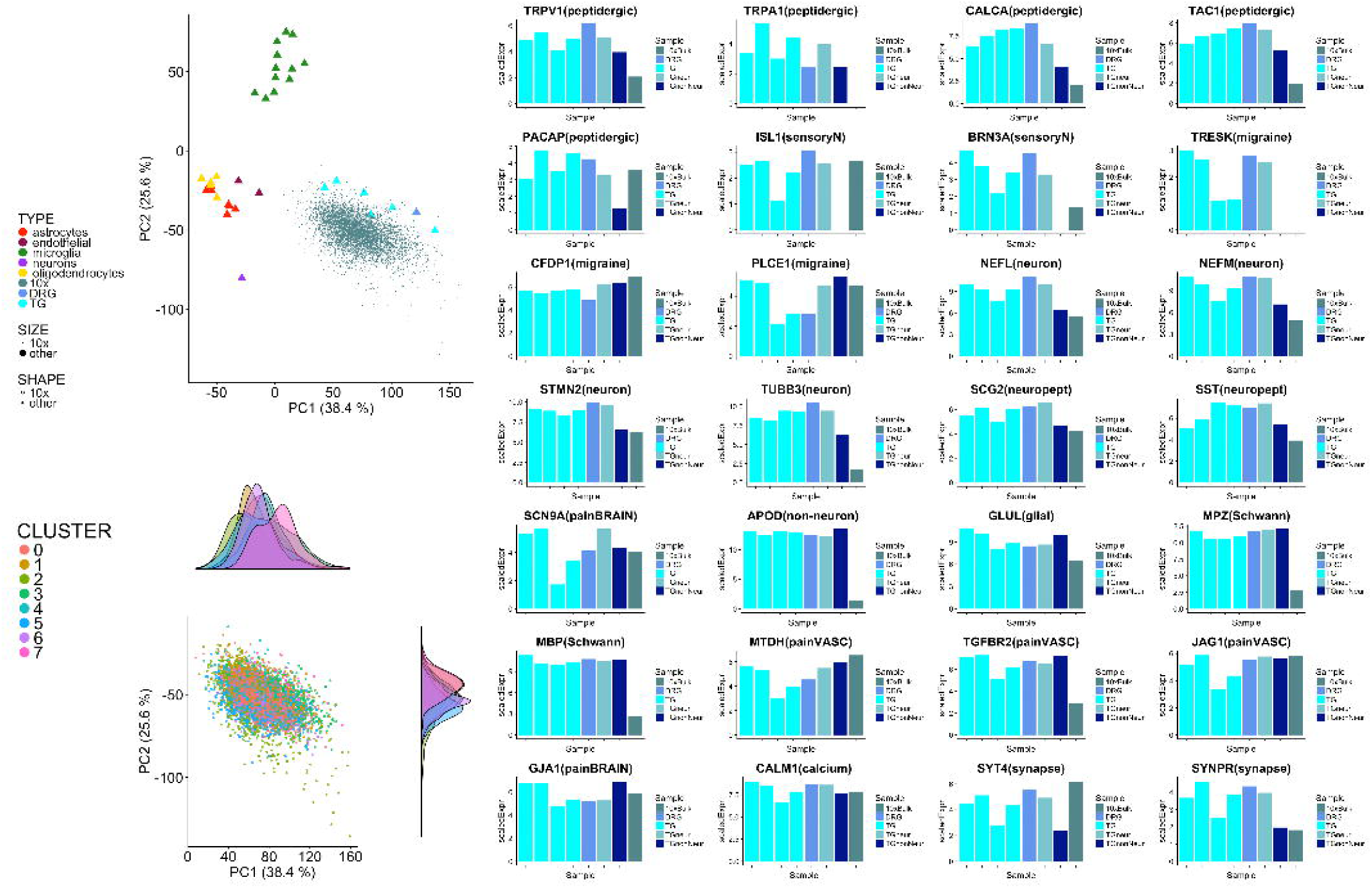
10x single cell analysis confirms peptidergic-like sensory neuron identity for all cells. (Left panel, top) Cell type identity space built through PCA of external RNAseq brain cell types. When projected onto this space, iPSC-derived nociceptor single cells cluster near the external TG and DRG samples. (Left Panel, bottom) iPSC-derived nociceptor single cells projected on the same space do not separate by the cell clusters identified Seurat (see Methods). (Right panel) Gene expression levels for given genes within the combined single cells (pseudo-bulk) obtained by merging all reads from all single cells, and for external TG and DRG samples. Gene expression is scaled on housekeeping genes. The functional classification for each gene is reported within parentheses based on information derived from ^27^. TG and DRG samples are from ^28^. TG “neuron” and TG “non-neuron” are from (LaPaglia et al., 2017).

The peptidergic identity of our replated TG nociceptors was confirmed by the expression of *TRPV1*, *CGRP* (*CALCA*), *TAC1* and *PACAP* similar to as observed in post-mortem human TG samples as previously described (LaPaglia et al., 2018). Despite TRPA1-agonist induced calcium flux was observed in these cultures, we did not detect the expression of *TRPA1* in the analysis, which perhaps could be attributed to low number of reads detected for this gene. These nociceptors also expressed sensory neuronal markers including *ISL1*, *BRN3A*, markers indicative of pain vasculature such as *MTDH*, *TGFBR2*, *JAG1*, synaptic markers *SYT4*, *SYNPR* and importantly, glial marker *GLUL* and Schwann cell marker *MPZ* (Fig. 5) which could perhaps be the key cell types facilitating the maturation of these replated TG cultures to release CGRP.

### Astrocytic co-culture of placodal isolated TG cultures promotes peptidergic nociceptive features

The single cell transcriptomic analysis on the replated peptidergic TG nociceptor cultures revealed the expression of glial markers, and it is widely recognized that astrocytes help support neuronal survival and maturation (Allen et al., 2012). We therefore tested whether glial co-culture could change the fate of our isolated TG placodal nociceptors from non-peptidergic to peptidergic fate. We observed that isolated TG placodal nociceptors co-cultured with primary rat astrocytes now expressed CGRP as confirmed by CGRP/TUJ1 co-expression in 66.78% of the cells whilst GFAP expressing astrocytes showed no CGRP expression (Fig. 6a). These co-cultures also exhibited 5-fold increase in basal CGRP release, compared to those cultured in the absence of astrocytes (Fig. 6b), although we did not observe inflammatory soup evoked release. Analysis by calcium imaging further confirmed functional competence in these neuron-astrocyte co-cultures, where approximately 60% of the nociceptors responded to 10µM capsaicin and no such response was seen in placodal isolated neurons cultured in the absence of astrocytes (Fig. 6c). Hence, this data suggests that the neuronal-astrocyte interactions aids in fate determination with the induction of CGRP expression.

**Figure 6:**
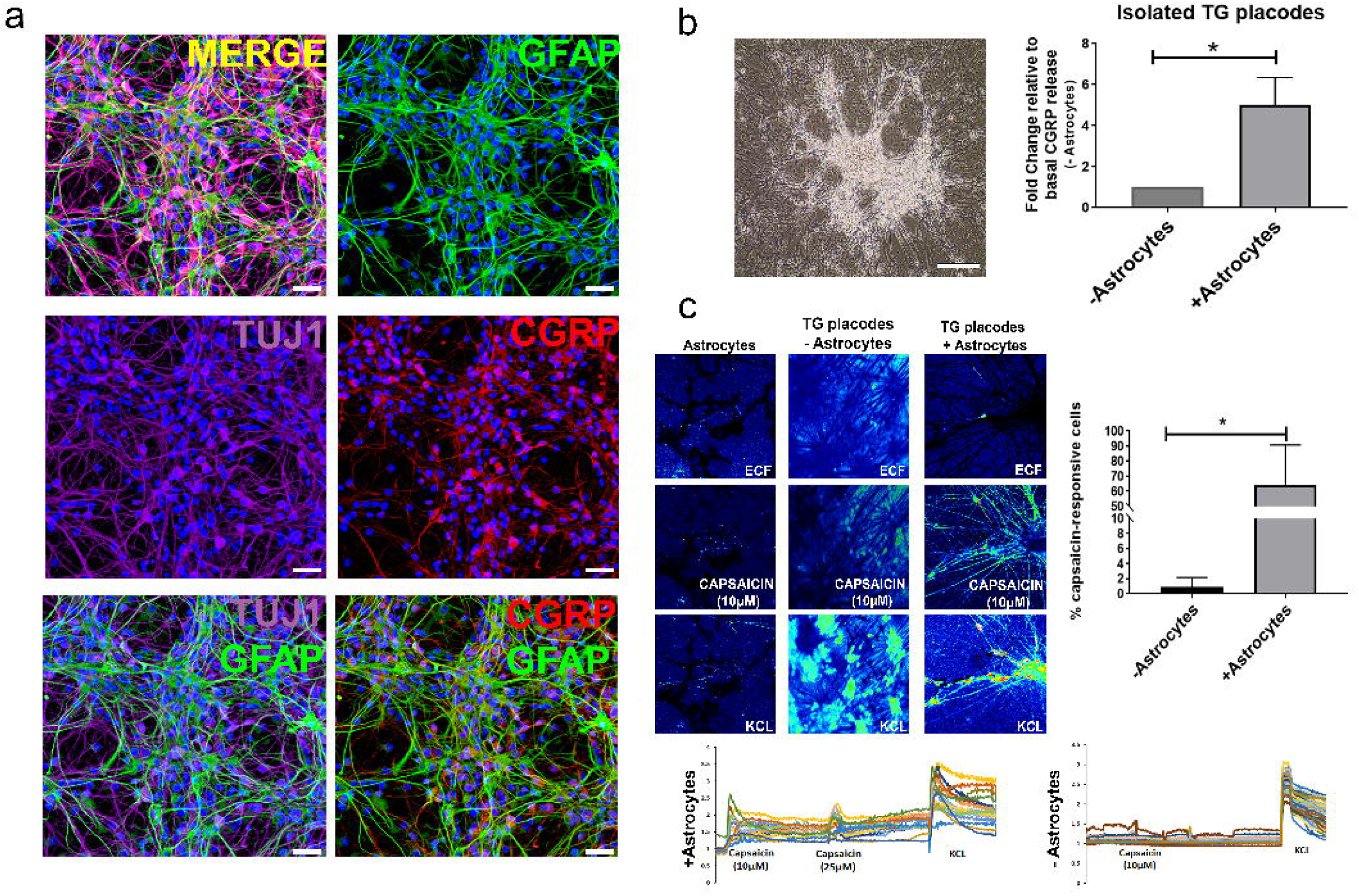
Co-culture of isolated TG placodes with astrocytes induces peptidergic nociceptive fate. **a)** Immunocytochemistry reveals peptidergic identity of TG placodes-astrocyte co-cultures indicated by GFAP-expressing astrocytes and neurons expressing both CGRP and TUJ1, scale bar 25µM. **b)** (Left panel) Brightfield image of neuron-astrocyte co-cultures, scale bar 10µM. (Right panel) Astrocyte co-cultures facilitate basal CGRP release compared to TG placodes cultured in the absence of astrocytes, (n = 3 separate dishes per iPSC line differentiation per condition, fold change quantified relative to basal release, p<0.05). **c)** Functional analysis via calcium imaging suggests increased sensitivity to capsaicin in placode-astrocyte co-cultures similar to replated cultures (n = 3 dishes from three independent differentiation rounds, with or without astrocytes, p<0.05).

## Discussion

The diversification of nociceptors into sub-classes is a multi-step process likely involving early patterning of neuronal progenitors followed by postmitotic differentiation towards a mature nociceptor subtype specific phenotype, governed by both intrinsic signals which determine their differentiation potential and the surrounding niche that influence the population size of nociceptor subtypes (Barres et al., 2015). Peptidergic neurons gain Runx1^+^/TrkA^+^/cMet^+^ identity whereas the non-peptidergic acquire Runx1^+^/TrkA^+^/Ret^+^ expression profile. As development progresses and functional maturation occurs through the perinatal and postnatal period, the peptidergic nociceptors lose Runx1, giving them a cMet^+^/TrkA^+^/Cgrp^+^ identity and the non-peptidergic neurons continue to express Ret, a receptor for a glial-derived neurotrophic factor (GDNF), thereby exhibiting a Runx1^+^/Ret^+^ expression profile (Gascon et al., 2010; Liu and Ma, 2011). Hence, the mechanism controlling nociceptor fate specification is attributed to intrinsic determinants such as functional expression of cMet, as well as to extrinsic factors including the balance of NGF, HGF and GDNF signaling which may be required for the persistence or extinction of Runx1 expression (Gascon et al., 2010).

We found in TG differentiation, that following placodal induction, the sensory neurons markers ISL1 and BRN3A were expressed, along with the nociceptor marker TrkA. We observed expression of RUNX1 at early stages which then increased along with expression of RET confirming a non-peptidergic nociceptor identity. A recently described comprehensive unbiased RNA-seq study of 622 mouse neurons (Usoskin et al., 2015) has brought to light the plethora of nociceptor subtypes that exist, at least based upon transcriptomic identity. The functional relevance of these many subtypes remains to be established. The TG placodal-isolated nociceptors comprised a large population of TRPA1^+^/TRPV1^-^/TRKA^+^/RET^+^/CGRP^-^ cells which are similar to the NP1 category from the RNAseq classification and a minority population of TRPA1^+^/TRPV1^+^/TRKA^+^/RET^+^/CGRP^-^ cells which match the non-peptidergic NP2/NP3 category. Interestingly we found that unlike neural crest derived non-peptidergic neurons from human iPSC, our TG placode derived cells did not respond to ATP, normally mediated by P2X3 receptors in sensory neurons. However an examination of rat TG sensory neurons indicated only a minority of IB4^+^ non-peptidergic neurons express P2X3, whilst in DRG IB4^+^ sensory neurons, 60-70% expressed P2X3 (Ambalavanar et al., 2005). Our findings are therefore consistent with a TG lineage of non-peptidergic nociceptors.

Peptidergic nociceptors are an important nociceptor subtype in health and disease including inflammatory and neuropathic pain and migraine (Lukacs et al., 2017), and a human *in vitro* model of these nociceptors would help advance disease modelling and therapy development for many pain-related disorders. We now report for the first time, the highly efficient generation of human iPSC-derived functional model of TG peptidergic nociceptors. Almost all the peptidergic nociceptors, expressed TRPV1, aligning with the peptidergic PEP1 category of mouse neurons (Usoskin et al., 2015) and were responsive to capsaicin. TRPV1^+^ peptidergic nociceptors are considered to mediate noxious thermal pain behaviors and consistently no cells responded to menthol, indicating an absence of functional cold sensor, TRPM8. PEP1 (CGRP+) and TRPM8 being two separate populations is fitting with the human TG sequencing data (Yang et al., 2022a).

A substantial proportion of peptidergic nociceptors expressed TRPA1 (Cavanaugh et al.; Le Pichon and Chesler, 2014; Ro et al., 2009). In mouse models, Trpv1 is expressed widely expressed in DRG neurons initially and then becomes restricted to peptidergic nociceptors. Distribution of TRPV1 in rat (Price and Flores, 2007) and human (Jung et al., 2023; Shiers et al., 2020, 2021; Tavares-Ferreira et al., 2022; Yang et al., 2022b) nociceptors appear to be broader, blurring the non-peptidergic and peptidergic boundary. We found that our iPSC differentiated nociceptors expressed functional TRPV1 in both non-peptidergic and peptidergic subtypes although the different nociceptor subpopulations had differential responsiveness to capsaicin (Demartini et al., 2017; Durham and Masterson, 2013; Hoffmann et al., 2013). The non-peptidergic nociceptors expressing TRPV1 only responded to 25µM capsaicin whilst peptidergic nociceptors were sensitive to 10µM capsaicin, as has been reported for mouse TG peptidergic nociceptors (Hoffmann et al., 2013). Similarly, we have also found that TRPA1 can be expressed in both peptidergic and non-peptidergic nociceptors.

The CALCA-GFP iPSC reporter line revealed that CGRP gene expression increases early in the TG differentiation protocol and is clearly present at the cranial placode stage before becoming rapidly extinguished after placodal isolation. The decline in CGRP expression corresponded to the decrease in cMET expression in these cultures, consistent with the previous studies indicating the importance of cMET in the production of peptidergic nociceptors (Gascon et al., 2010). Although some studies have indicated a low overall level of CGRP in sensory neurons during the pre-natal period, earlier studies have demonstrated that TG neurons innervating the cerebrovasculature are enriched for CGRP expression (O’connor and Van Der Kooy, 1988). TG neurons innervate peripheral targets by E14-15 and are expressing CGRP a few days later (Tsai et al., 1989). Hence our *in vitro* TG neurons may be analogous to those TG neurons that supply to the cerebral blood vessels, with early expression of CGRP. Our single cell transcriptomic analysis also confirmed the similarity of TG peptidergic nociceptors with the previously published data on human TG (LaPaglia et al., 2018). Whilst there were apparent distinct sub-clusters present when examining the iPSC nociceptors alone, once projected onto the cell type-specific gene expression space, cell clusters were not defined by varying cell identity probably due to the very low and noisy expression of nociceptor marker genes at the single cell level.

Given the early expression of cMET and CGRP, it seemed that with the appropriate signals, the TG cultures could adopt a peptidergic fate. This was confirmed when we either did not isolate the placode or replated the whole culture after the placode had formed at a lower density and found the overwhelming majority of neurons were peptidergic nociceptors. This suggests that other cells in the culture outside of the cranial placode are providing an extrinsic signal for peptidergic fate specification perhaps akin to target derived signals *in vivo*. Remarkably we found that astrocytes may able to provide much of the extrinsic signaling required to generate peptidergic nociceptors from the cranial placode. Astrocytes are widely recognized to support neuronal function, promote neuronal survival and maturation by releasing trophic factors including NGF and BDNF during development (Allen and Barres, 2009; Allen et al., 2012; Seth and Koul, 2008). We also demonstrated CGRP expression in 65.8% of cells the TG placodal isolated nociceptors which otherwise exhibit only non-peptidergic features in the absence of astrocytes. Whilst we observed a basal release of CGRP with astrocyte co-culture, we were unable to evoke release with inflammatory soup, suggesting that other factors may be required for evoked release.

In rodent models of migraine, application of inflammatory soup to the dura leads to activation of TG neurons, increased CGRP release into the jugular vein (Hoffmann et al., 2009) and increased CGRP expression in the TG (Lukács et al., 2015). In our cultures, exposure of the TG peptidergic nociceptors to inflammatory soup potently led to increased release of CGRP. Moreover, we found that the migraine provocants PACAP-38 but not GTN, was able to evoke CGRP release. We speculate that CGRP release in our *in vitro* nociceptors is mediated via cAMP pathway since we also observed CGRP release with FK but not with cGMP-pathway agonists including SNAP, GTN and 8-bromo-cGMP (De Mey and Vanhoutte, 2014; Miller and Megson, 2007) (Fig. 3f). Sumatriptan is a highly effective acute migraine therapeutic drug and was developed to target the 5-HT1 receptor subtypes (5-HT_1B_, 5-HT_1D,_ 5-HT_1F_) on blood vessels to induce vasoconstriction (Ahn and Basbaum, 2007). The vascular changes in migraine are now not considered primary and instead there is considerable focus on CGRP, based upon increased CGRP levels during migraine attacks and the success of clinical trials of CGRP receptor antagonists and more recently the monoclonal antibodies against CGRP or the CGRP receptor (Goadsby et al., 2017; Oliveira et al., 2024; Silberstein et al., 2017). The 5-HT_1B/1D/1F_ receptors have been shown to localize on the presynaptic terminals of TG neurons (Ahn and Basbaum, 2007) and sumatriptan has been shown to directly inhibit the stimulated but not the basal CGRP release from cultured rat TG neurons (Durham and Russo, 1999). Importantly we found that sumatriptan was able to prevent the stimulated release of CGRP but did not affect basal CGRP release. We also observed inhibitory effect of topiramate on stimulated CGRP release. The ability of migraine provocants to induce CGRP release and migraine treatments to reduce CGRP release supports the importance of CGRP in migraine pathophysiology and also illustrates the potential value of iPSC-peptidergic nociceptors as a disease model and basis of a phenotypic screening assay.

Future work will investigate the interaction of maturing nociceptors with the adjacent cells including glia to decipher CGRP release mechanism, as it is clearly apparent that replating placodes with the surrounding niche promotes generation of functional TG peptidergic nociceptors, whilst isolating the placodes in the absence of astrocytes leads to the induction of non-peptidergic nociceptors. The knowledge gained in this study paves way to generate a more *in vivo*-like ganglionic composition of non-peptidergic and peptidergic nociceptors to identify specific signaling cues and cross-talk between the different sensory neuronal subtypes. This will enable translational studies to improve drug discovery in pain disorders that continue to face significant unmet medical need.

### Online Methods Human iPSC culture

Human iPSC lines (AH017-7, SBAD-03-01, SBAD-02-01) were derived from fibroblasts of healthy individuals after informed consent, using CytoTune-iPS Sendai Reprogramming Kit (Life Technologies). The iPSCs were derived as part of the IMI/EU sponsored StemBANCC consortium and underwent full quality control (QC) analysis including FACS for pluripotency markers NANOG/TRA-1-60, SNP karyotyping, post-thaw viability and mycoplasma testing and were considered suitable for experiments as they were mycoplasma negative, expressed pluripotency markers and genomically stable. Cell lines were cultured in feeder-free system on Matrigel™ (Corning) and were fed with mTESR1™ media (STEMCELL Technologies). Cultures were passaged every 5-7 days or when 75-95% confluent, using EDTA (Life Technologies) and split at 1:4 ratio with medium changed every day. iPSCs were thawed post-QC freeze and used for experiments with five passages.

### Generation of CGRP-GFP targeting plasmid vector

To construct pUC57CGRP-GFP-T2A-Puro for expression of GFP reporter, CGRP promoter sequence -2053 to +91 (Genebank ID: AC090835.7, bases 45451-47599) was synthesized by ThermoFisher Scientific. The Splice acceptor-T2A-puro was amplified by PCR from a plasmid using primer pair 5′ACGCGTCACTCTCGAGGGAGAGGGCAGAGGAAGTCTT CTAAC3′ and 5′-ACGCGTCACTCTCGAGCCATAGAGCCCACCGCATCCCC-3′ (XhoI site underlined). The PCR product was ligated with the XhoI sites of the vector pUC57CGRP-GFP. The resulting plasmid pUC57CGRP-GFP-SA-T2A-Puro, consisted of Zinc Finger site (ZFN) for AAVS1 locus.

### Transfection of iPSCs with targeting vector and Zinc Finger Nuclease (ZFN) plasmids

StemBANCC iPSC line SBAD03-01 was selected for targeting as it was derived from a healthy donor, showed a stable karyotype and efficiently differentiated towards all three lineages (data not shown). SBAD03-01 cells were harvested using TrypLE (Gibco) and 300.000 cells/well of a 12 well plate were co-transfected with two plasmids: one expressing Zinc Finger Nucleases (ZFN) and the other containing the CALCA promoter, fluorescent reporter and a drug-resistance cassette using Lipofectamine LTX (ThermoFisher Scientific). We used a targeting plasmid vector, pUC57CALCA-GFP-T2A-Puro, in which the last GFP coding codon is fused in frame with a T2A sequence followed by puromycin. Both ends of the insert are loxP-flanked (floxed) in order to remove it if needed. Transfected cells were passaged once confluent and selection with puromycin (0.2µg/ml) was performed after 24 hours. When puromycin-resistant colonies expanded, single cell cloning was performed through serial dilution in 384 well plates to identify pure clones.

### Junction PCR and droplet digital PCR (ddPCR)

Junction PCR and ddPCR analysis identified multiple correctly targeted clones without random transgene integration from a total of 300+ puromycin-resistant clones screened. Junction PCR was performed using primers pairs from both 5’ end and 3’ end of the insert to identify clones with insert at correct location. DNA from clones was isolated using Modified Gitschier Buffer (MGB) and PCR was performed using Phusion high fidelity Taq pol (ThermoFisher Scientific). For ddPCR analysis, droplets were generated using QX200 droplet generator and FAM/Hex labelled probes. PCR products were then analyzed by QX200 droplet reader and QuantaSoft software.

The targeting efficiency (7%) was comparable to the efficiencies observed with TALENs and CRISPR/Cas9 using similar targeting strategies (Hockemeyer et al., 2009, 2011). CGRP-GFP clones 1-4, 5-17 & 5-40 were identified as single copy insertion and 1-8, 1-14 & 1-38 were identified as two copy insertion based on ddPCR data. For our study, CGRP-GFP clone C1-4 was employed.

### Differentiation of human iPSCs towards TG nociceptors

Directed differentiation of iPSCs towards TG nociceptors was performed as previously described (Dincer et al., 2013). Briefly, cells were lifted as single cells in accutase (Stem Cell technologies) and plated at 60.000 cells/cm^2^ in the presence of 10µM Y-27632 (Abcam) and 10ng/ml bFGF (Peprotech) in MEF-conditioned media. Upon 90-95% confluency, neural induction was initiated using 250ng/ml Noggin (R&D Systems) and 10µM SB431542 (Tocris) for 3 days in basal media composed of DMEM-F12, Glutamax, 20% Knockout Serum Replacement (Life Technologies). From day 4, cells were cultured only in the presence of 10µM SB431542 and from day 5 onwards, N2B27 media (B27 supplement without vitamin A, Life Technologies) was added in increments of 25%, 50%, 75% every 2 days, for 13-17 days leading to the formation of placodes. Next, three different maturation strategies were adopted. First, only the placodes were manually isolated and plated on matrigel up to days 55-65 in maturation media composed of N2B27, 20ng/ml BDNF (Peprotech), 0.2mM Ascorbic acid (Sigma) and 2µM Forskolin (Sigma). Second, the TG placodes were not isolated and the maturation medium was added from day 15 onwards. Third, day 15 cultures were lifted using accutase (placodes and the surrounding cells) and they were replated on matrigel at 200.000/cm^2^ cell density in maturation medium followed by downstream analysis day 55 onwards. All three iPSC lines underwent three independent differentiations towards TG nociceptors.

### Immunocytochemistry

Samples were fixed in 4% paraformaldehyde (Sigma) for 20 minutes at room temperature (RT). They were then permeabilized for eight minutes in 0.1% Triton X-100 (Sigma) diluted in 1X phosphate buffered saline (PBS), washed in 1X PBS for five minutes and blocked for one hour in 1% BSA, 0.05% Tween-20 in 1X PBS at RT. Subsequently, cells were incubated in primary antibodies (diluted in blocking solution) at 4°C overnight, washed three times in 1X PBS for five minutes and treated with secondary antibodies (diluted in blocking solution) for one hour at RT. Next, samples were washed again three time in 1X PBS for five minutes, incubated with DAPI for 10 minutes and mounted on glass slides using Dako Mounting Fluorescence medium (Dako). Primary antibodies used were rabbit anti-PAX6 (1:200, Biolegend), goat anti-SIX1 (1:100, SCBT), rabbit anti-CASPASE3 (1:200, ), goat anti-RUNX1 (1:200, SCBT), rabbit anti-cMET (1:200, Abcam), goat anti-PERIPHERIN (1:100, SCBT), rabbit anti-ISLET1 (1:200, Abcam), rabbit anti-BRN3A (1:100, Abcam), mouse anti-TUJ1 (1:1000, Biolegend), rabbit anti-SYNAPTOPHYSIN (1:100, Abcam), rabbit anti-TRPV1 (1:100, Alomone Labs), rabbit anti-5HT_1D_ (1:100, Alomone labs), mouse anti-CGRP (1:100, Abcam), rabbit anti-TRKA (1:100, Alomone Labs), rabbit-anti RET (1:200, Alomone Labs), rabbit anti-NEUN (1:500, Milipore), chicken anti-GFAP (Abcam) and mouse anti-TUJ1 (Abcam). Secondary antibodies used were donkey anti-goat Alexa488, donkey anti-rabbit Alexa594, goat anti-rabbit Alexa488, goat anti-mouse Alexa594, goat anti-mouse Alexa488 and goat anti-rabbit Alexa594. All secondary antibodies were from ThermoFisher Scientific and were used at 1:500 dilution. All images were acquired by confocal microscopy using Zeiss 880 inverted microscope (Zeiss).

### Quantitative real-time PCR (qRT-PCR)

Total RNA was isolated using TRIzol and RNeasy Mini kit (Qiagen). Reverse transcription was performed on 300ng RNA per sample using SuperScript First-Strand Synthesis kit (Thermofisher Scientific). The cDNA was diluted 1:5 for each sample and qRT-PCR was performed using SybrGreen PCR Master Mix (ThermoFisher Scientific) and Applied Biosystems 7500 Fast Real-Time PCR machine. Housekeeping genes included *GAPDH* and *βACTIN* and genes of interest used were *RUNX1*, *cMET* and *CGRP*. All samples were analysed as biological triplicates (n = 3 for each sample). Negative controls included both water (no cDNA) and no RT (RNA) samples. The CT values were normalized against housekeeping genes and fold change in gene expression was analyzed using ΔΔCT method. Statistical significance was determined using one way-ANOVA and p<0.05 was considered significant.

### CGRP ELISA

Medium was collected from the different culture conditions, centrifuged at 200g for 5 minutes and supernatants were frozen at -80°C in multiple aliquots to avoid freeze-thaw cycles. Briefly, basal media, 80mM KCl stimulated-media (4-hour exposure) and inflammatory soup^21^, 2µM FK, 1µM PACAP-38, 1mM SNAP, 1µM 8-bromo-cGMP-stimulated media and 1µM GTN (Sigma) were collected following 24 hours exposure (n = 3 separate dishes per iPSC line for each treatment). For drug-inhibition assay, media was collected from the same dishes that were first pre-treated with 10µM sumatriptan or 100µM topiramate (Sigma) for one hour, followed by exposure to I.S. supplemented with either of the drugs (n = 3 separate dishes per iPSC line per treatment). CGRP ELISA was conducted using Human CGRP Enzyme Immunoassay Kit as per manufacturer’s instruction (Bertin Pharmacy/Bioquote). Plates were read on a Wallac Victor 1420 Multilabel counter at 405nm for 0.1s and data was analyzed on GraphPad Prism 6 software. Statistical significance was determined using one way-ANOVA and p<0.05 was considered significant.

### Calcium imaging

Mature nociceptors from all the three culture methods (placode isolation, no isolation and replating) were loaded with 2µM Fura2-AM and 80µM Pluronic acid (ThermoFisher Scientific) and incubated at 37°C for one hour. They were then washed 1X PBS and fed with in-house formulated extracellular fluid (ECF, composed of 145mM NaCl, 5mM KCl, 10mM HEPES, 10mM D-Glucose, 2mM CaCl_2_, 1mM MgCl_2_, pH 7.4). For imaging, excitation filters BP340/30 and BP387/15 were used to capture images every 2 seconds and the ratiometric 340/380 calculations were performed by subtracting the background. Baseline imaging was performed for 30 seconds in ECF followed by 30 second exposure to the agonists with five-minute ECF wash between each agonist perfusion. ATP (10µM) was applied for 30 seconds, followed by capsaicin (10µM, 25µM) and finally KCl (50mM) or the cells were exposed sequentially to menthol (100nM), mustard oil (250µM), capsaicin (10µM, 25µM) and KCl (50mM).

### Flow Cytometry

Cells were treated with accutase for 10 minutes, centrifuged at 400g for 5 minutes and the cell pellet was re-suspended in 2% PFA (Alfa Aesar) diluted in FACS buffer (1% FBS, 10µg/ml human serum albumin and 0.01% Azide made up in 1X PBS) for 10 minutes at RT. They were then permeabilized in 1ml ice cold 100% methanol for 30 minutes at -20°C or stored up to several weeks for later use. Next, approximately 500,000 cells were harvested, spun at 2000rpm for three minutes, followed by wash in FACS buffer and split into two for each antibody combination. Cells were incubated in primary antibody for 1 hour at RT, washed in FACS buffer, incubated with secondary antibodies for 45 minutes at RT and washed and re-suspended in FACS buffer for analysis. Primary and secondary antibodies used were the same as previously described for immunocytochemistry including NEUN, TRPV1 and CGRP.

### Electrophysiology

Human iPSC-derived nociceptors were plated directly onto the Ibidi dishes (Thistle Scientific) for functional characterization. Individual ibidis were placed in a recording chamber mounted onto the stage of an upright microscope. For targeted whole-cell recordings, we used a 40x water immersion objective and IR-DIC optics. Cells were continuously superfused with oxygenated aCSF (95% O_2_/5% CO_2_) containing 130mm NaCl, 25mm NaHCO_3_, 2.5mm KCl, 1.25mm NaH_2_PO_4_, 2mm CaCl_2_, 2mm MgCl_2_ and 10mm glucose. Patch-clamp electrodes (4–7 MΩ) were filled with an intracellular solution containing 120mm K-gluconate, 10mm KCl, 10mm HEPES, 4mm MgATP, 0.3mm NaGTP and 10mm Na-phosphocreatine. Recordings were obtained using a Multiclamp 700B amplifier and digitized at 10–20 kHz using Digidata 1550 acquisition board.

### Sample preparation for 10x genomics

iPSC-TG nociceptors were washed in PBS with 0.04% BSA and re-suspended at a concentration of ∼1500cells/μl before capturing single cells in droplets on Chromium 10x Genomics platform. Library generation for 10x Genomics v2 chemistry was performed following the Chromium Single Cell 3ʹ Reagents Kits User Guide: CG00052 Rev B. Quantification of cDNA was performed using Qubit dsDNA HS Assay Kit (Life Technologies Q32851). Quantification of library construction was performed using Qubit dsDNA HS Assay Kit (Life Technologies Q32851) and high-sensitivity DNA tapestation (Agilent. 5067-5584). Libraries were sequenced on Illumina HiSeq4000 platform to achieve an average of 42,000 reads per cell.

### Alignment, barcode assignment and UMI counting

We used the pipelines “mkfastq” and “count” from the Cell Ranger Single-Cell Software Suite (https://support.10xgenomics.com/single-cell-gene-expression/software/pipelines/) to demultiplex raw base call files into FASTQ files, and to perform alignment, barcode counting and UMI counting. Multiple sequencing runs (two HiSeq4000 lanes) of the same library were combined by the “count” pipeline. Barcodes and UMIs were filtered using default settings to generate gene-barcode matrix.

### Clustering analysis

We used Seurat R package (Butler et al., 2018) for QC filtering and data analysis. Only genes with at least one UMI count detected in at least one cell were used. Only cells with less than 10% of sequenced mitochondrial genes and more than 200 expressed genes were kept. This left 12257 cells and 25541 genes for analyses, with approximately ∼2,000 genes expressed in each cell. UMI normalization was performed using the function “NormalizeData” that consisted in dividing UMI counts by the total expression in each cell, followed by multiplication with a scale factor (10,000 by default), and by log-transforming the UMI counts. The top 2,000 most variable genes were identified using the function “FindVariableGenes” with parameters x.low.cutoff = 0.0125, x.high.cutoff = 3, y.cutoff = 0.3. We regressed out cell-cell depth variation in gene expression using the number of detected UMIs with the function “ScaleData”. The function “FindClusters” was used to perform a graph-based clustering approach on the normalised and scaled dataset with parameters reduction.type = "pca", dims.use = 1:5, resolution = 0.3. To confirm the identified clusters, we used an unsupervised clustering method for single cell RNAseq data called SC3 (Kiselev et al., 2017). We found that five of the eight clusters identified by the two methods shared more than 50% of the cells while three were less robust.

### Cell type gene expression PCA space

We merged RNA-Seq data from purified human brain cell types including neurons, astrocytes, oligodendrocytes and endothelial cells available at http://www.brainrnaseq.org (Zhang et al., 2016), from isolated microglia during surgery of cortical regions in patients with epilepsy (Gosselin et al., 2017), from human trigeminal ganglia (TG) and dorsal root ganglia (DRG) (Flegel et al., 2015) and from human TG (LaPaglia et al., 2018). PCA (scaled and centered) was performed on the merged dataset and the internal 10x single cells were projected onto this space using the “predict” R function.

## Acknowledgements

The research leading to these results has received support from the Innovative Medicines Initiative Joint Undertaking under grant agreement n° 115439, resources of which are composed of financial contribution from the European Union’s Seventh Framework Programme (FP7/2007-2013) and EFPIA companies’ in-kind contribution. This research has also been supported by the NIHR Oxford Biomedical Research Centre. This publication reflects only the author’s views and neither the IMI JU nor EFPIA nor the European Commission are liable for any use that may be made of the information contained therein. We would also like to thank Prof. Dave Bennett, Dr. Alex Clark and Dr. Gregory Weir for their help with calcium imaging set-up and analysis as well as critical reading of the manuscript.

## Author Contributions

Conceptualization, G.D. and Z.C.; Methodology, G.D., S.C., S.K.G, M.M, R.H.; Software, C.W; Validation, G.D., S.C., C.H., P.P., T.L., C.H., D.C., K.A., XL; Formal Analysis, G.D., V.V., C.W.; Investigation, G.D., S.C.; Resources, T.L., R.B., H.S., I.J.O., Z.C.; Data curation, V.V, C.W.; Writing – Original Draft, G.D.; Writing – Review & Editing, G.D., Z.C; Visualization, G.D., T.L., Z.C.; Supervision, Z.C.; Project Administration, G.D., S.C., Z.C; Funding Acquisition, Z.C.

## Supplementary Figures

**Supplementary Figure 1: Optimization of TG differentiation protocol.** TG nociceptors generated in the presence of matrigel and FK combination **a)** reduces cell death as shown by CASPASE-3 expression and **b)** are functional and have a physiological resting membrane potential of -60mV (n = 12 cells per iPSC line, p<0.05).

**Supplementary Figure 2: RUNX1 and cMET expression profiling during DRG nociceptor differentiation to identify peptidergic and non-peptidergic population. a)** RUNX1 expression increases while cMET expression decreases as differentiation progresses, scale bar 10µm. Dashed box reveals small population that has RUNX1^+^/cMET^-^ non- peptidergic identity at day 40, scale bar 25µm. **b)** Gene expression profiling by qRT-PCR demonstrates significant increase in non-peptidergic *RUNX1* marker while peptidergic markers *cMET* and *CGRP* are significantly downregulated (p<0.05, n = 3 biological replicates).

**Supplementary Figure 3: Schematic showing the detail map of CGRP-GFP-T2A-Puro insert**. This was used in the generation of CGRP-GFP reporter line.

**Supplementary Figure 4: ObLiGaRe Mediated Targeting of the AAVS1 Locus through drug selection. a)** Schematics of the targeting strategy. The presence of the ZFNs and targeting plasmid vector, results in the insertion of pCGRP-GFP-T2A-Puro cassette at AAVS1 locus in human chromosome 19. In the targeting schematics here, colored boxes are various segments of the insert that is flanked by LoxP sequence. **b)** The PCR primers (GFP-F + R) used for genotyping for identifying correctly targeted clones. Six of the correctly targeted clones (clone 1-8, 1-14, 1-38, 1-4, 5-17 & 5-40) were selected for further characterization & validation. **c)** ddPCR analysis indicating insert copy number. Internal puromycin probe was used for analyzing the clones, showed zero insert copy number for control iPSC, two copies for clones 1-8, 1-14 & 1-38 and one copy for clones 1-4, 5-17 & 5-40. Blue bar represents total events xnumber for puromycin probe and the green bar represents total number of events for reference gene AP3B1. Aqua sample had no template DNA.

**Supplementary Figure 5: Validation of TG peptidergic nociceptors derived upon non-placodal isolation replating strategy**. These neurons expressed nociceptive markers (ISL1, PERIPHERIN), neuronal marker TUJ1, synaptic marker SYNAPTOPHYSIN and peptidergic marker CGRP, scale bars 25µm, 50µm, 25µm (L-R).

**Supplementary Figure 6: Electrophysiological analysis of TG peptidergic nociceptors derived upon non-placodal isolation replating strategy**. The somas of differentiated neurons were targeted by DIC for whole cell recordings as illustrated in the figure (n = 15 cells per iPSC line). Example traces of current clamp recordings of repetitive firing and occasional spontaneous activity in iPSC-derived nociceptors. Voltage clamp recordings show large inward Na currents that are TTX sensitive. Peptidergic neurons display repetitive firing in response to depolarization and spontaneous activity indicating functional neuron/maturation (left panel)

**Supplementary Figure 7: Co-expression of 5HT_1D_ receptor and CGRP.** Analysis by immunocytochemistry in non**-**placodal isolated replated TG peptidergic nociceptor cultures, scale bar 10µm.

