## Supplementary figures and images for "A highly efficient method to differentiate CGRP-expressing peptidergic nociceptors from human induced pluripotent stem cells"

### Supplementary Figures

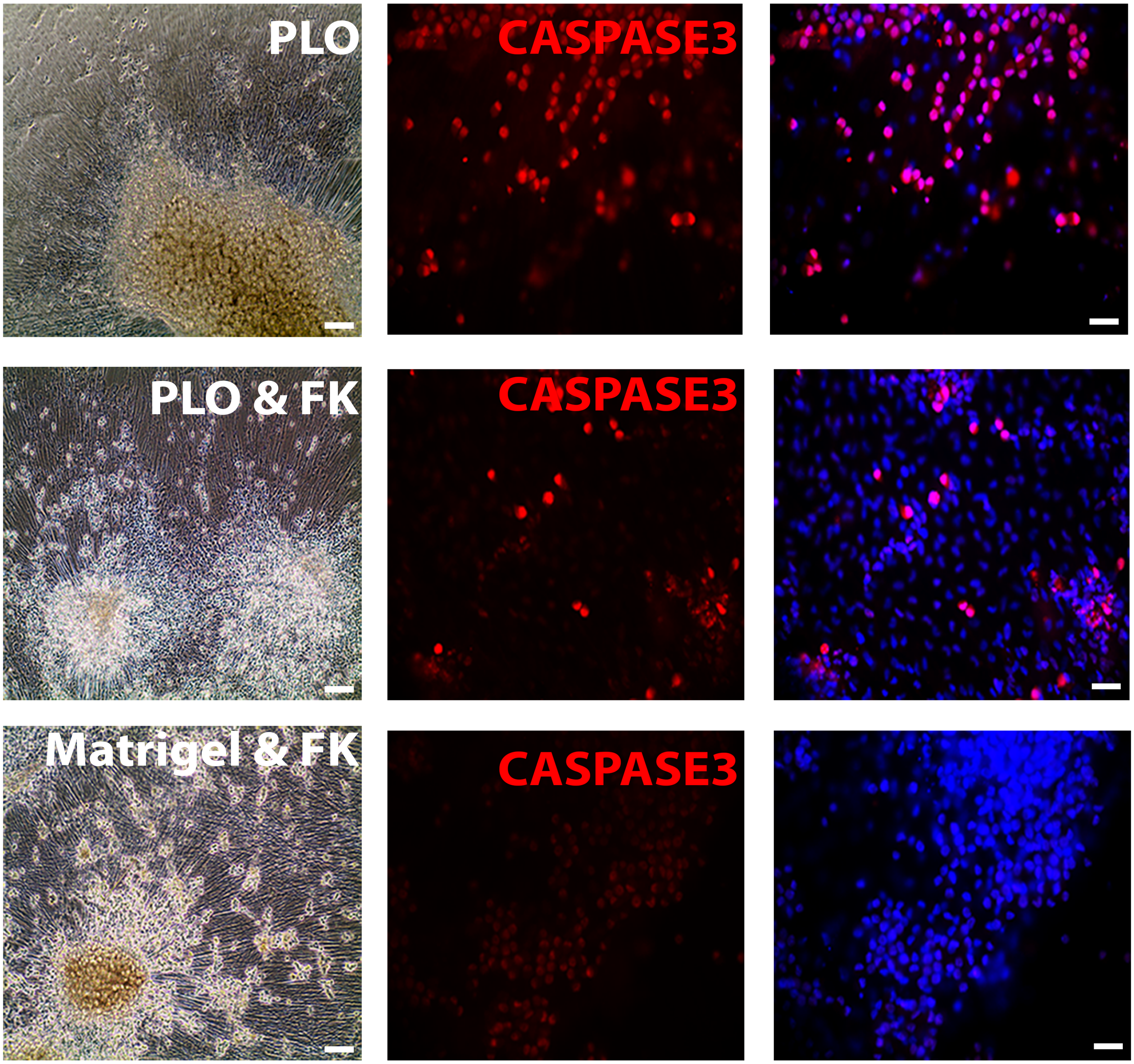

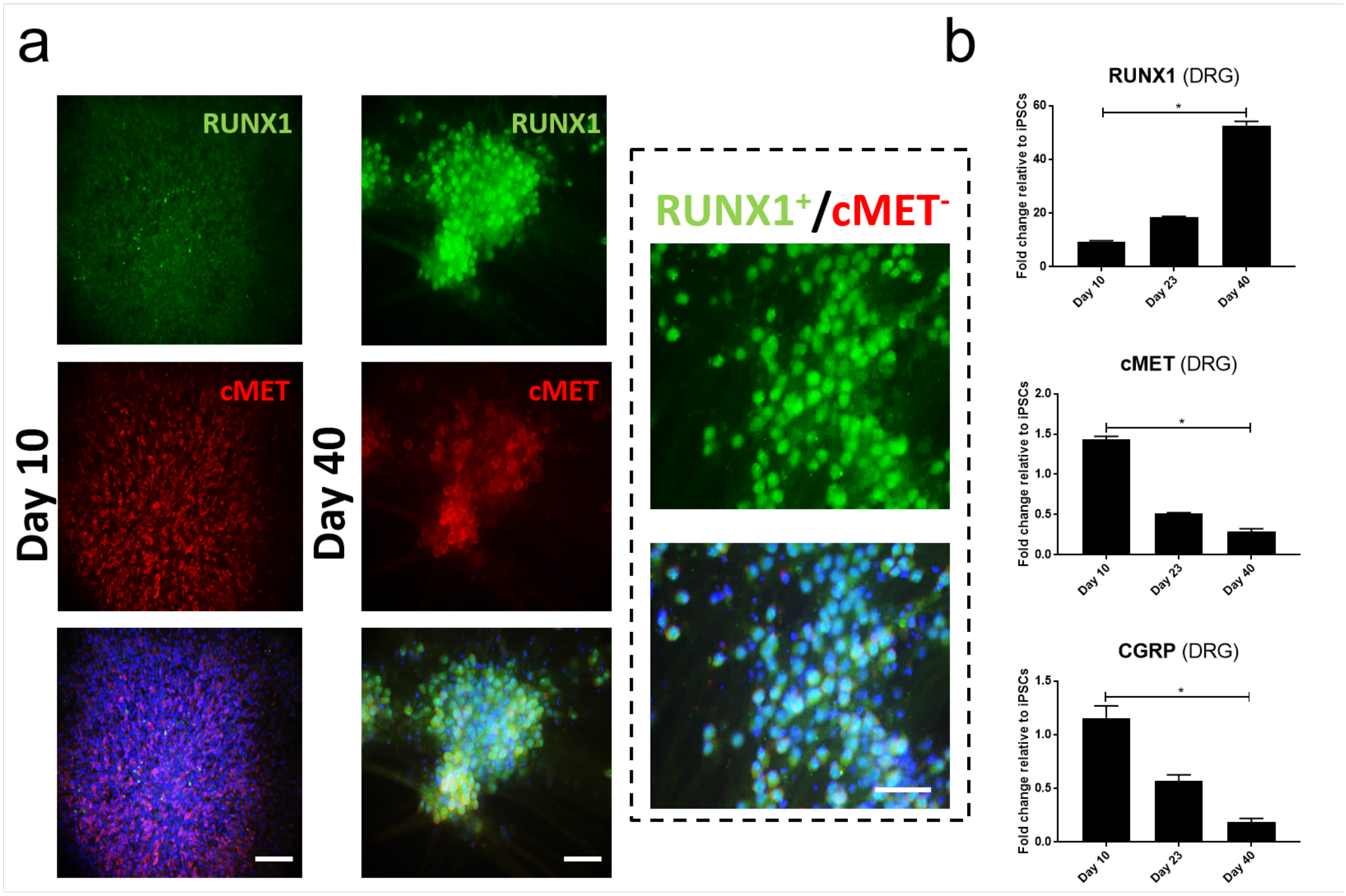

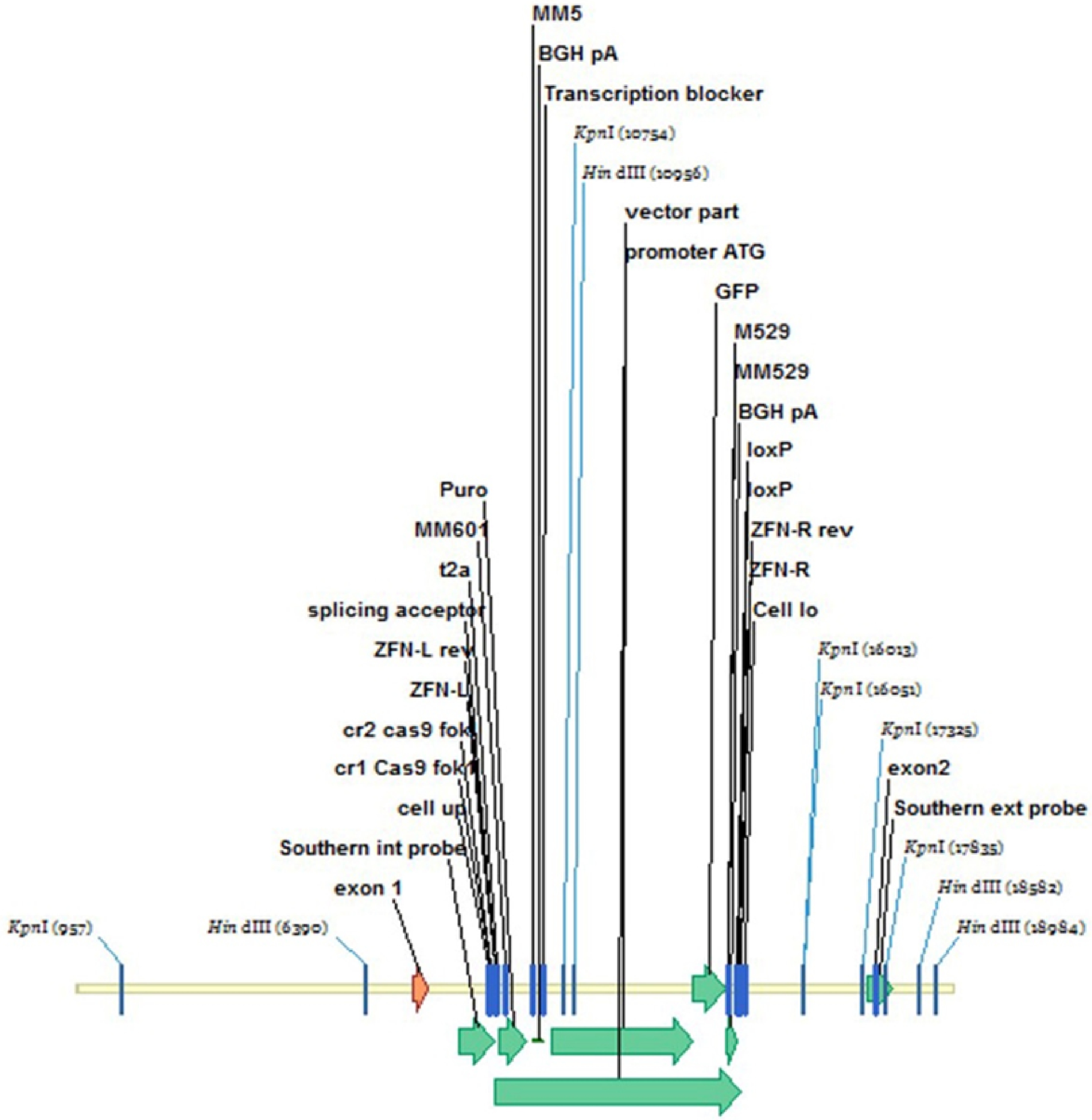


hAAVS1:CGRP-GFP
(19335bp)

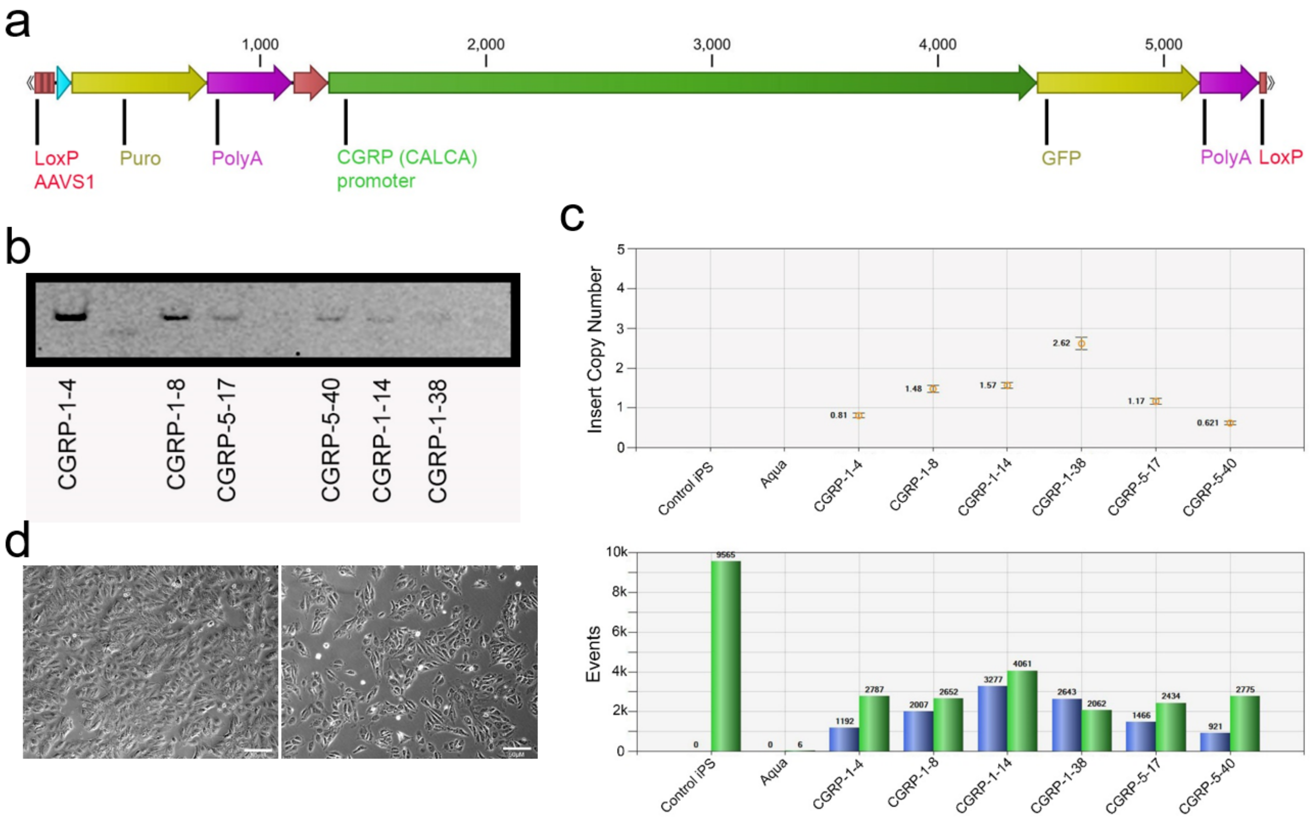

#### Mature peptidergic nociceptor population

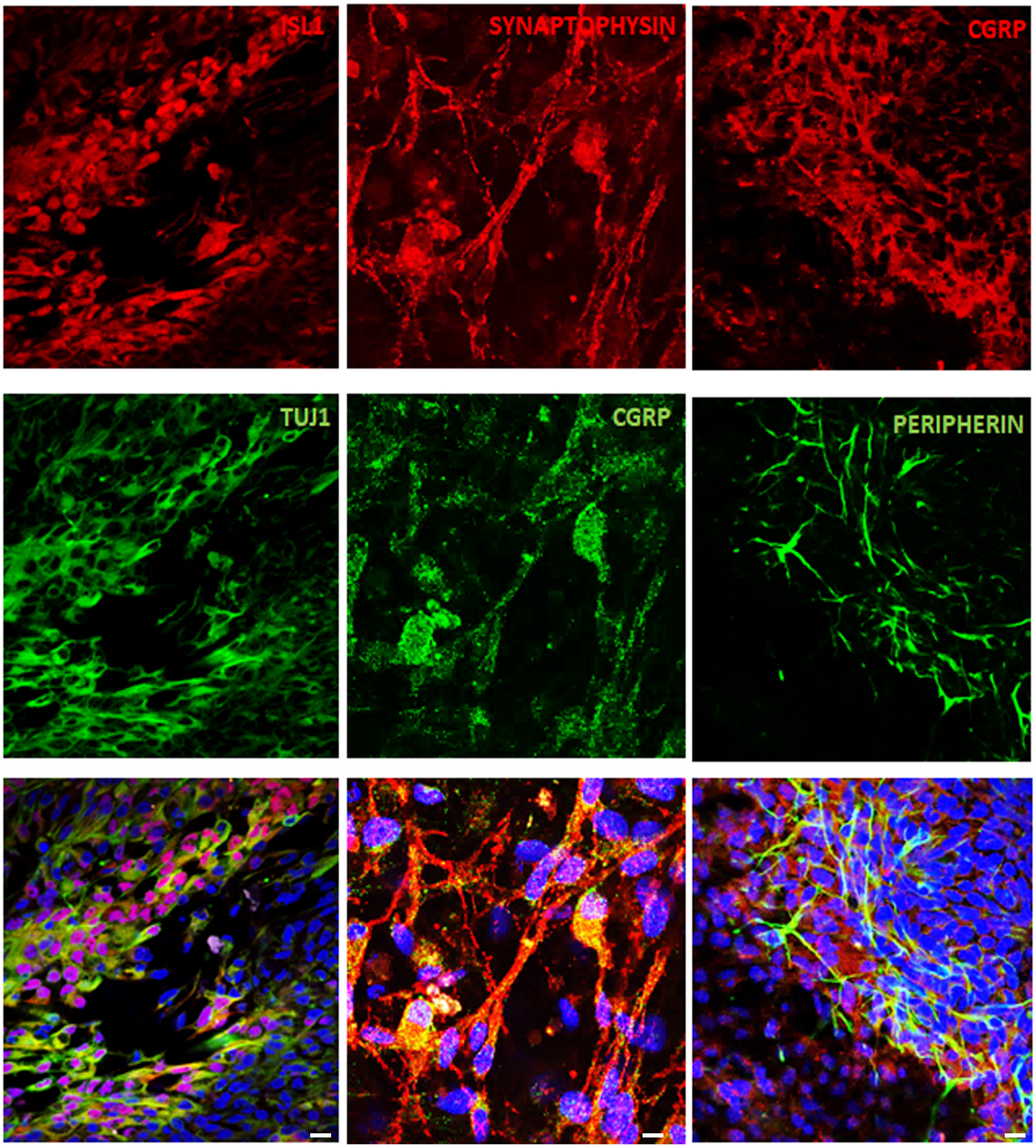

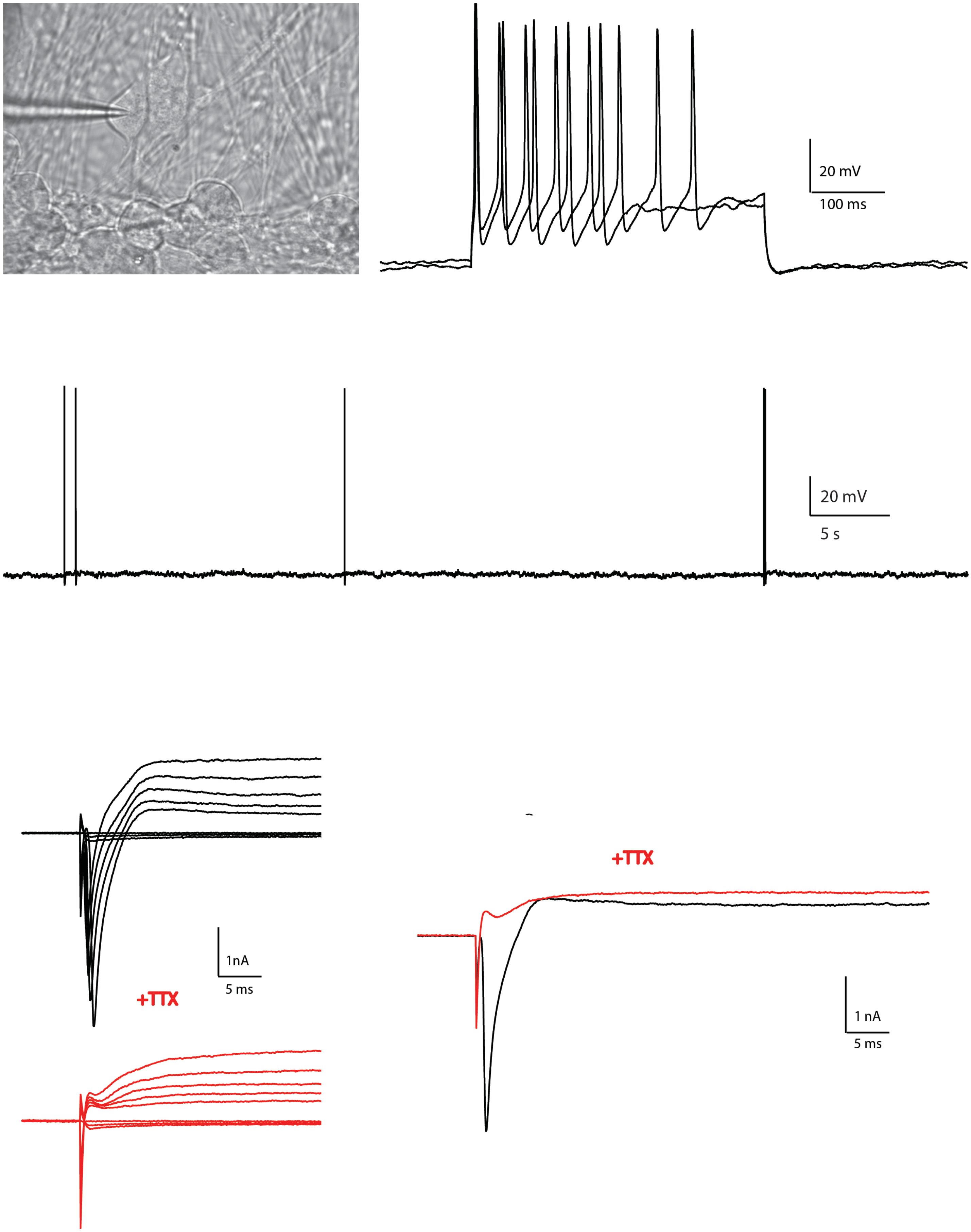

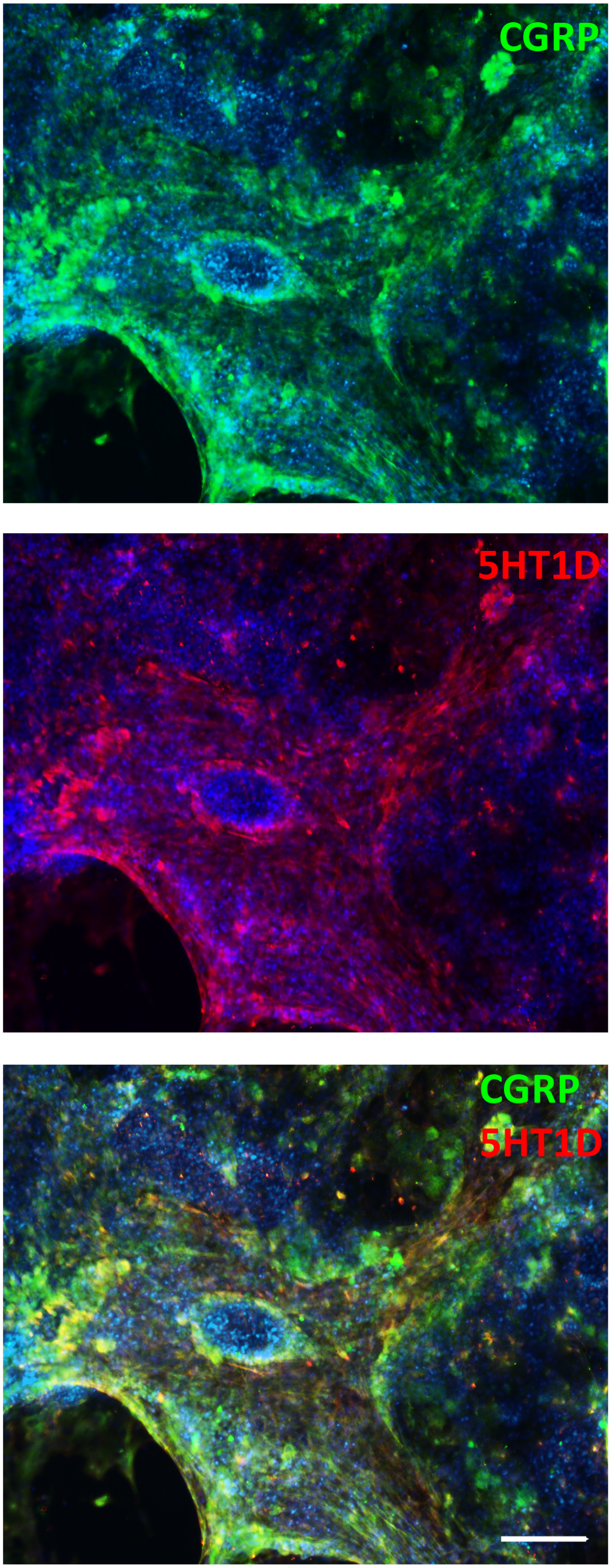
